# Long-lasting impact of starvation experience on fly activity and place preference

**DOI:** 10.1101/2020.12.10.419622

**Authors:** Deepthi Mahishi, Tilman Triphan, Ricarda Hesse, Wolf Huetteroth

**Affiliations:** University of Leipzig, Faculty of Life Sciences, Department of Genetics, Talstrasse 33, 04103 Leipzig, Germany

**Keywords:** *Drosophila*, feeding, foraging, place preference, tracking

## Abstract

Animal behaviours are demonstrably governed by sensory stimulation, previous experience and internal states like hunger. With increasing hunger, priorities shift towards foraging and feeding. During foraging, flies are known to employ efficient path integration strategies. However, general long-term activity patterns for both hungry and satiated flies in conditions of foraging remain to be better understood. Similarly, little is known about how chronic contact chemosensory stimulation affects locomotion. To address these questions, we have developed a novel, simplistic fly activity tracking setup – the Panopticon. Using a 3D-printed Petri dish inset, our assay allows recording of walking behaviour, of several flies in parallel, with all arena surfaces covered by a uniform substrate layer. We tested two constellations of providing food: i) in single patches, and ii) omnipresent within the substrate layer. Fly tracking is done with FIJI, further assessment, analysis and presentation is done with a custom-built MATLAB analysis framework. We find that starvation history leads to a long-lasting reduction in locomotion, as well as a delayed place preference for food patches not driven by immediate hunger motivation.

## Introduction

Flies show hunger-motivated ranging or foraging walks to find food; they also show explorative walks (or local searching behaviour) after food encounter (Dethier, 1957; Bell et al., 1985; Bell, 1990; Corrales-Carvajal et al., 2016; Kim and Dickinson, 2017; Murata et al., 2017; Hughson et al., 2018; Mahishi and Huetteroth, 2019), and much has been achieved in identifying the circuits and dynamics involved in this behaviour (Corfas et al., 2019; Lin et al., 2019; Moreira et al., 2019; Sayin et al., 2019; Seidenbecher et al., 2020). Most of these studies either used hunger-motivated behaviour to focus on the underlying navigational strategy of the flies, or they focused on exploration-exploitation trade-offs under different motivational settings.

Interestingly, both hedonic and caloric value of the food source can influence explorative walks. The perceived sweetness after ingestion correlates to the duration and path length of explorative walking (Murata et al., 2017), and protein-sated flies venture further away from a yeast patch than protein-deprived flies (Corrales-Carvajal et al., 2016). As satiation levels drop with ongoing post-prandial energy expenditure, finding food is becoming increasingly important. An exploring fly constantly assesses palatability with its tarsal chemosensors, supported by occasional proboscis extension (Mahishi and Huetteroth, 2019). But how much does a fly explore when nutritional homeostasis can be achieved anywhere?

What are the locomotion dynamics independent of food search? To overcome distorting foraging locomotion, driven by constantly changing hunger levels, we provide an arena with omnipresent food. Although these previous studies imply foraging-independent explorative walking in sated flies (Bell et al., 1985; Bell, 1990; Murata et al., 2017), no study exists to our knowledge that studied prolonged intrinsic walking behaviour on a homogenous food substrate, where any locomotion motivated by food-seeking can be ruled out.

It is also not well understood how palatability and satiation affect walking activity beyond the first 3-120 min after food interaction; food-related responses can exert their physiological and behavioural effects on longer timescales. Larval diet composition impacts adult food choice (Davies et al., 2018), and preference of a caloric diet over an equally palatable alternative is only established after several hours (Dus et al., 2011; Stafford et al., 2012). Similarly, a dietary imbalance between palatability and nutritional content is leading to sustained physiological changes much later (Wang et al., 2016, 2017; Musso et al., 2017; Park et al., 2017; May et al., 2019). Apart from few exceptions (Martin, 2004; Meunier et al., 2007), most existing fly locomotion studies either examine short periods at high temporal resolution (Kim and Dickinson, 2017; Murata et al., 2017; Brockmann et al., 2018; Hughson et al., 2018; Landayan et al., 2018), or sample for short recurring time windows to cover longer periods (Green, 1964a, 1964b; Connolly, 1966a; Barwell et al., 2020). Automated circadian studies provide both temporal resolution and timespan, but focus on changes in rhythmicity (Guo et al., 2016; Pegoraro et al., 2020), and rarely include location preferences or locomotion in a nutritional context (Donelson et al., 2012; Dreyer et al., 2019).

We use uninterrupted video tracking (1 Hz) of starved and fed flies to compare locomotion activity and location probability for over 24 h in two conditions: i) a foraging setting with a single food patch, or ii) with homogenous food substrate on all surfaces. Our assay comes with a data pipeline from recording to analysis utilizing custom-written camera recording software, FIJI-based tracking, and MATLAB-based data analysis, various sanity check functions and visualization.

## Material & Methods

### Animals

All experiments were performed with 2-5 day old male Oregon^R^ flies, maintained at the University of Leipzig at 25 °C and 60 % humidity on a 14:10 LD cycle on standard fly food. Starved animals were kept for 22 - 24 h in empty vials with added wet tissue, fed flies were allowed to feed *ad libitum* on normal fly food until testing.

### Panopticon assay

Cold anesthetised fed and starved male flies were alternately placed in individual sectors of the Panopticon and then transferred onto the imaging rig (Fig. 1A), which is located inside a climate chamber (not shown) to maintain constant levels of 60 % humidity and a temperature of 25 °C. The Panopticon consists of a 3D-printed arena (.stl file; Renkforce RF1000: Material PLA white), which is inserted in the lid of a standard plastic Petri dish (85 mm, Greiner) partially filled with substrate (1 % agarose or 1 % agarose containing 200 mM sucrose), separating it in eight sectors (5.5 cm² each) (Fig. 1B). During insertion of the plastic arena into still viscous substrate we assured homogenous coating of all inner walls before solidification. For foraging experiments, small Eppendorf lids were used to create individual food containers in each arena (food patches, 0.2 cm²). The 1 % agarose with 200 mM sucrose in these food patches was levelled with the surrounding 1 % agarose to avoid confounding effects of negative geotaxis on place preference (Robie et al., 2010). The Panopticon was closed with another inverted Petri dish lid with a layer of substrate to provide equal surface texture on all sides (Fig. 1B). Data collection was started as soon as all flies regained walking ability (within 1 min of transfer). Images (1024 × 1024 px) were recorded with a camera (Basler acA1300-200uc) at 1 frame per second for 24 h.

**Figure 1:**
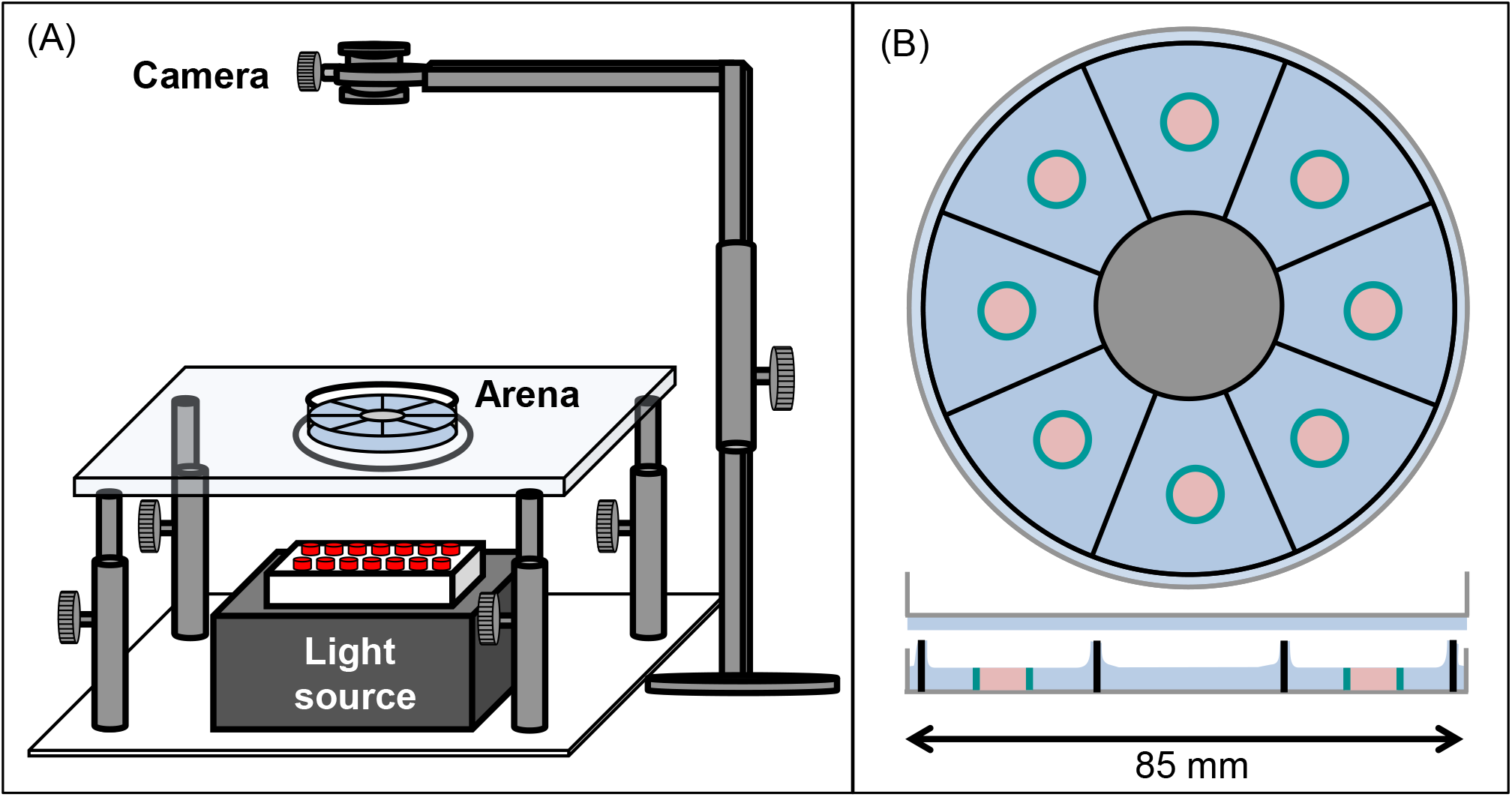
The Panopticon. **(A)** Entire recording setup for the Panopticon. The whole assay platform has an infrared light source placed in the bottom (grey box with red LEDs), on top of which is a height-adjustable transparent glass platform with a small circular plastic layer acting as as diffusor. The arena itself is placed on this platform, positioned precisely in line with the light source, the diffusor and the recording camera fitted on top with a stable holder. **(B)** Top view of Panopticon, sagittal view below. The arena consists of eight sectors, as indicated by sector-dividing walls of the 3D printed inset (black), which sits in substrate (blue) inside a Petri dish lid (gray). Another Petri dish lid with a layer of substrate closes the arena. Two configurations are used: i) food patch arena with 1 % agarose as substrate (depicted), which contains individual, centrally located food patch containers (teal blue) filled with 1 % agarose with 200 mM sucrose (pastel pink), or ii) omnipresent food arena without food patches, but 1 % agarose with 200 mM sucrose as substrate (not shown).

### Tracking and analysis

The recorded images were processed in batches using a custom-written, FIJI-based macro. Object information was extracted and saved as csv files. Results were then further handled using a data analysis script written in MATLAB 2018a, which provides various sanity check functions and visualization. Some plots make use of the “shadedErrorBar” function (Campbell, 2020). Activity and place preference plots are either aligned according to start time (1 h plots) or time of day (24 h plots). To exclude movements caused by camera noise we set a minimum threshold of 2 pixels (0.1 mm) and treated the fly as stationary in this case. Regions of interest for place preference (food patches, same-sized virtual food patches) were created in MATLAB. For further details on analysis and plotting functions see Results and Code, and the Readme file available on Github (https://github.com/dmahishi/Panopticon.git). Fast and efficient data handling allows for tracking and analyses of 24 h data (recorded at 1 Hz) in about 4 h on a regular desktop PC (Ryzen 3 3200G, 16 GB RAM, Intel SSD 660p 512GB).

### flyPAD

Food sip measurements were performed on the flyPAD (Itskov et al., 2014). Animals were fed or starved as before, anesthetized on ice and then transferred into individual flyPAD chambers with added wet tissue to allow for 24 h recordings. Data analysis took place with a customized version of the MATLAB script for the flyPAD (Itskov et al., 2014).

### Statistics

All statistical analyses were conducted using GraphPad Prism 8®. Multiple pairwise comparisons (paired t-tests) in case of activity and place preference plots and two-way ANOVA tests for displacement distribution and stop duration plots with Bonferroni-Dunn correction were performed.

## Results

### Paradigm and data pipeline for image based tracking of fly locomotion

We designed a novel, low-cost fly activity tracking setup, the Panopticon (Fig. 1). Walking behaviour of eight flies can be recorded simultaneously for up to 24 h. All arena walls and the top lid are covered in 1 % agarose (with or without 200 mM sucrose) to both provide equal surface texture on all sides and to maintain humidity levels and avoid desiccation, allowing continuous recordings for up to a week (data not shown). Experiments were performed in constant darkness (DD) with IR lighting (850 nm) to provide constant environmental conditions. It has been previously shown that DD is less disruptive to fly activity pattern than constant light (LL) (Green, 1964a).

We devised a complete data pipeline from recording to analysis (Fig. 2). Image recording utilizes custom-written camera recording software. Subsequently, the recorded images are analysed in batches using a custom-written, FIJI-based macro script. After background subtraction and pre-processing of the images objects are extracted and saved. The results are further analysed in a custom, MATLAB-based data analysis framework. Here, basic quality control and error checking functions are applied. At first, we calculated the rate of missed/failed detections, and experiments with an error rate of more than 5% were excluded from further analysis (Suppl. Fig. S1A). In the second step, we did a visual inspection of the walking traces (shown with a temporal colour code) to check for obvious detection errors (Suppl. Fig. S1B,C). As the last step we did a sector-wise plotting of particularly long frame-to-frame movement events that could be associated with potentially false-positive detections (Suppl. Fig. S1D). After this initial quality control, derived parameters like activity patterns and location probabilities are calculated.

**Figure 2:**
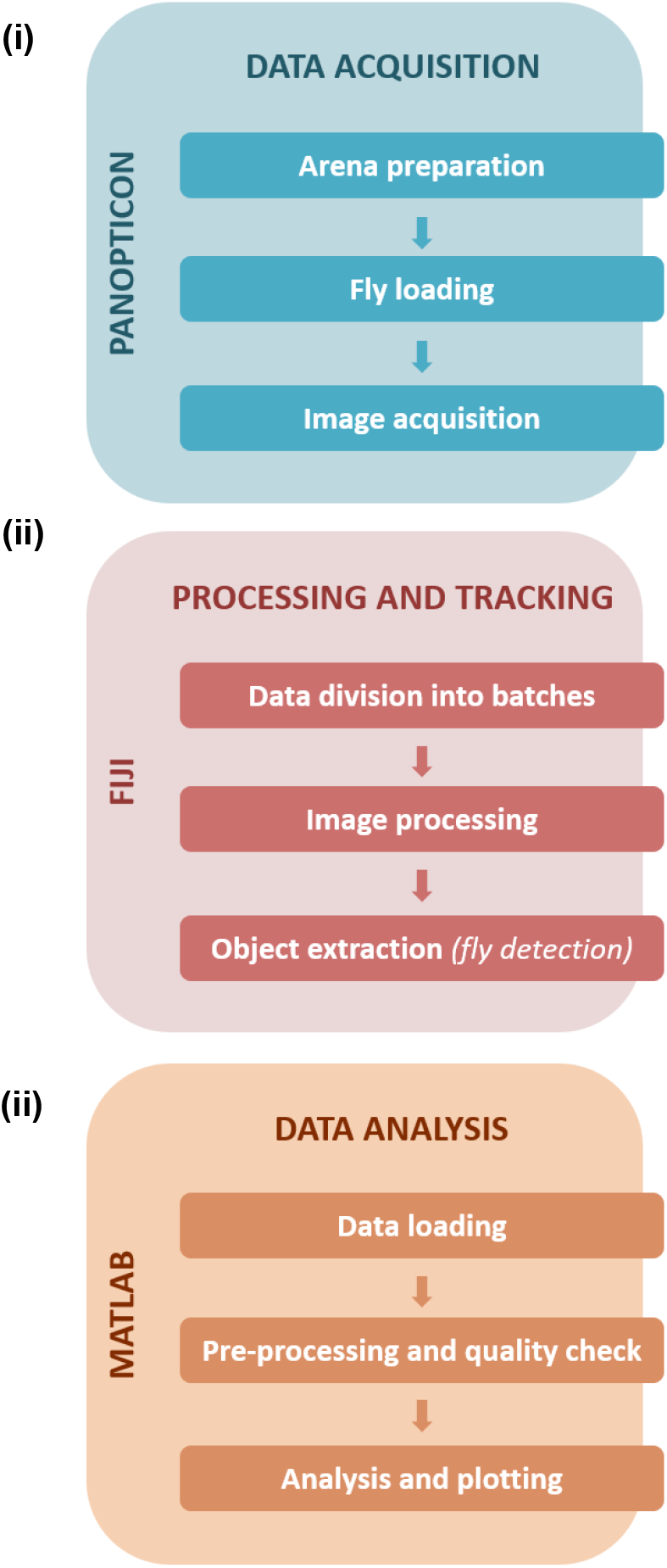
Data handling and workflow. Data acquisition (**i**) steps with the Panopticon involve preparation of the arena with substrate media, collection and loading of single flies in each of the sectors, followed by recording of fly activity. Further processing and tracking (**ii**) of recorded data (in JPEG format) is performed on FIJI to further divide collected frames into batches to facilitate efficient image processing and subsequent detection of fly positions. Data analysis (**iii**) was executed in MATLAB and consisted of pre-processing and quality checks followed by in-depth analysis and plotting of the tracked data (saved and loaded as3 .csv files).

### Starved flies appear sated within 30 min of food provision

First we assessed how hunger impacts locomotion activity during food search, and within which timescale the effect subsides after provision of food. Finding a food patch is well studied, and we know from existing data that locomotion increases with starvation level (Connolly, 1966b; Knoppien et al., 2000) and changes its dynamics after food patch encounter (Dethier, 1976; Corrales-Carvajal et al., 2016; Kim and Dickinson, 2017; Murata et al., 2017). Using the Panopticon with food patches, we find that both starved and fed flies exhibit increased activity for about 20-30 min in their new environment, as reported before (Soibam et al., 2012). This initial increased activity is significantly more pronounced in starved flies during the first 30 min (Fig. 3A), which correlates with compensatory overeating as soon as food is available (Carvalho et al., 2005). Flies need to stop to feed, and there is a reciprocal relation between the two behaviours (Mann et al., 2013). Accordingly, there is a significantly higher number of short duration stops (2-7 s) on the food patch during the first 30 min (Fig. 3B), which soon shift during the subsequent 30 min towards less frequent, longer breaks (Fig. 3C). At this point, both starved and fed flies reach equally low levels of activity (Fig. 3A). This pattern is reflected in sip numbers, as independently determined on the flyPAD, which provides a good estimate of actual food intake (Itskov et al., 2014). Starved flies exhibit a significantly higher sip number as fed flies during the first 20-30 min (Fig. 3D). After 30 min, sip numbers reach equal baseline levels (Fig. 3E), suggesting comparable satiation in both initially starved and fed flies. This is in accordance with previous studies that demonstrated reduced activity after a meal (Murphy et al., 2016).

**Figure 3:**
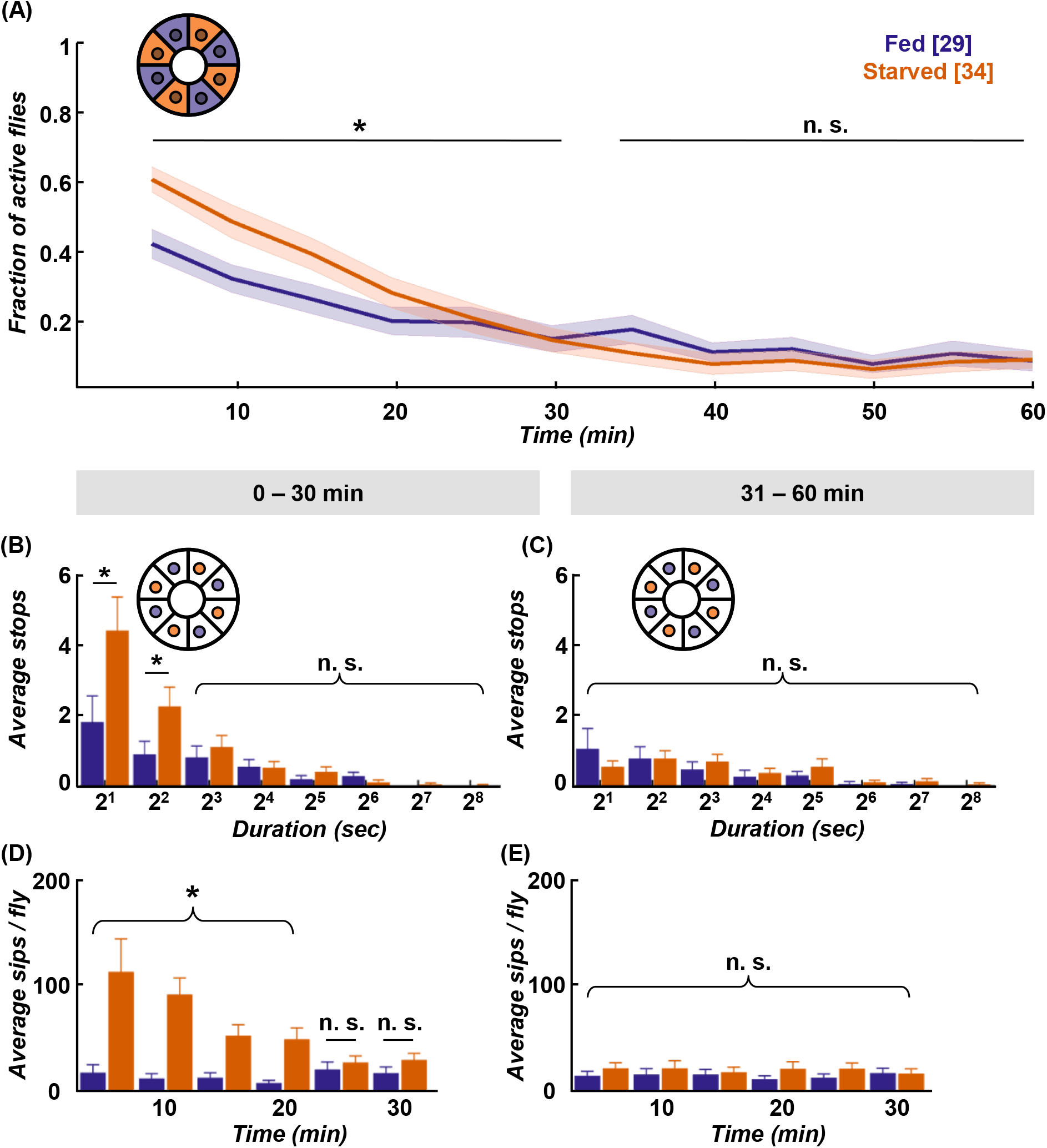
Starved flies show higher activity and food intake within first 30 min. **(A)** Both fed (deep purple) and starved (orange) flies exhibit raised activity on the food patch assay (inset schematic) during the first 30 min, with starved flies showing significantly higher activity than fed flies. Both groups reach baseline levels in the following 30 min. Data is presented as means ± s.e.m in 5 min bins. N values are given in brackets. **(B)** Initially starved flies make significantly more stops of 2–7 s duration than fed flies in the first 30 min, **(C)** but not in the next 30 min on the food patch (inset schematic). **(D)** Feeding rate, as measured on the flyPAD, indicates that initially starved flies show higher sip rate in the first 30 min than the fed flies, **(E)** but decreases to comparable sip rate in the next 30 min. * indicate significance levels following pairwise comparison or 2-way ANOVA tests (alpha = 0.05). Horizontal lines represent comparison between fed and pre-starved flies from a single group. Curly braces represent identical significance levels across multiple groups.

Taken together, our data shows that during the first 30 min starved flies exhibit increased activity, and an increased number of stops lasting between 2-7 s. These most likely reflect feeding bouts, which soon disappear when they presumably reach hunger motivation levels comparable to fed flies during the following 30 min.

### Starvation state affects 24 h walking activity and place preference in a food patch assay

Interestingly, despite the quick compensation in feeding motivation, we observe behavioural differences in locomotion between experimental groups in the long term. Fed flies exhibit a characteristic evening activity peak before the subjective night, which is missing in starved flies (Fig. 4A). This elevated evening activity in fed flies is accompanied by a significantly higher number of short-stop events in a representative 30 min time window (Fig. 4B). On the morning of the next subjective day, stop rates decreased to comparably low levels in both experimental groups (Fig. 4C). To see if this bias during the subjective evening is reflected in speed characteristics, we looked at displacement between frames as a proxy for velocity. However, displacement up to 2 mm/s on the food patch in the same time window (19:00-19:30) is not significantly different between fed flies and starved flies (Fig. 4D). On the subjective next morning (08:30-09:00), movement has slowed down equally in both groups (Fig. 4E). Whereas locomotive differences between initially starved and fed flies disappear on the subjective morning next day, another effect in positional preference becomes more pronounced; initially starved flies increasingly prefer to sit on or close to the food patch (Fig. 4F).

**Figure 4:**
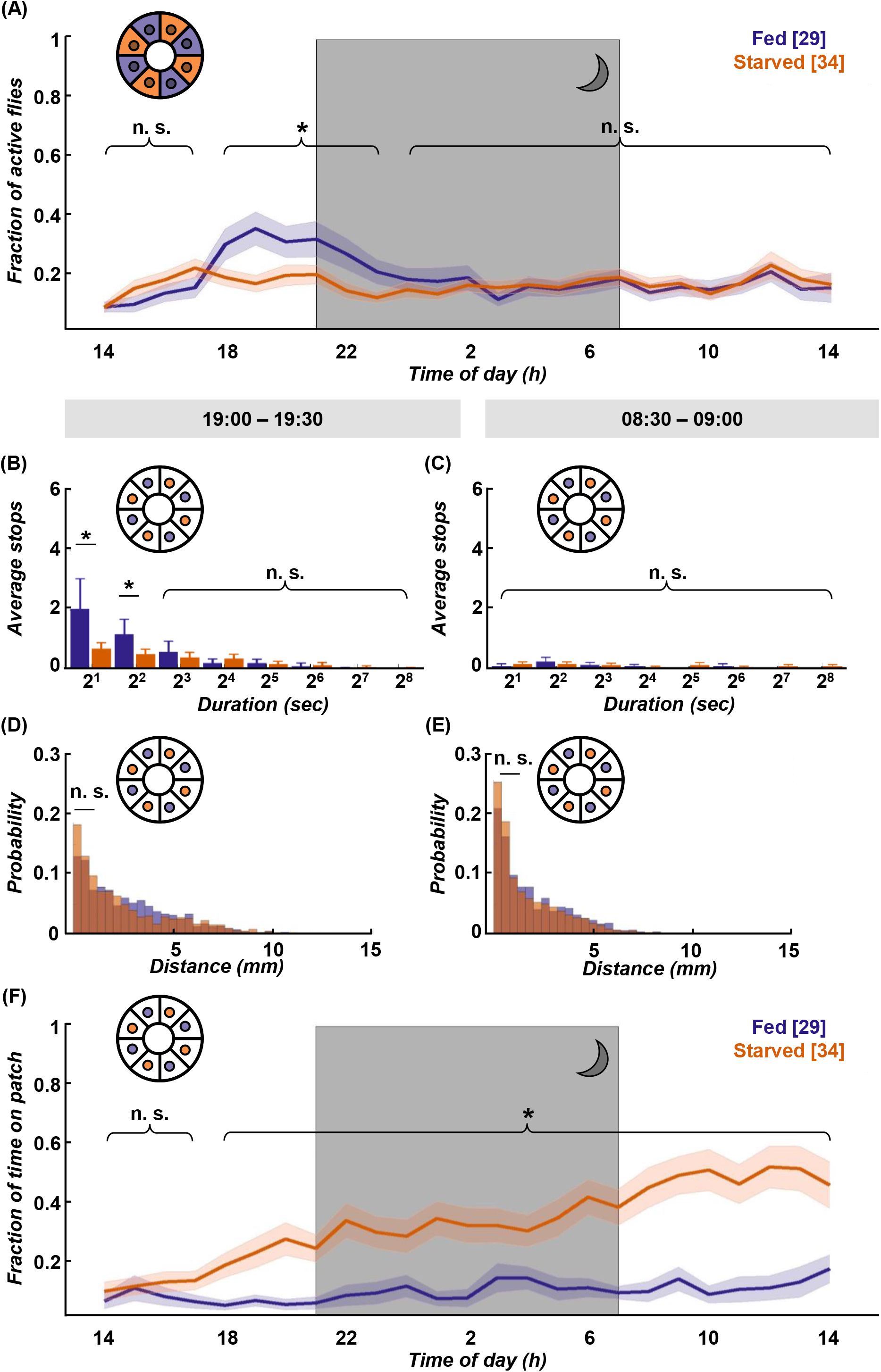
Locomotion activity and food patch preference over 24 h on food patch assay. **(A)** Although initial activity levels on the food patch arena (inset schematic) are comparable, pre-starved flies (orange, n = 34) continued to show significantly decreased activity levels during subjective evening and early night, as compared to fed flies (deep purple, n = 29). Subjective night indicated by grey-shaded area. There was no significant difference in activity across the rest of the 24 h. **(B)** Fed flies showed significantly increased numbers of stops of shorter duration (2-7s) on the food patch (inset schematic) than pre-starved flies during peak activity time window of 19:00-19:30, **(C)** whereas no significant difference in food patch stops was observed between 08:30-09:00 the next morning between the two groups. X-axis tick labels indicate duration of stops with a log2 scale, where 2^1^ = 2-3s, 2^2^ = 4-7s and so on. **(D)** On the food patch (inset schematic) there was no significant difference observed in short-distance moves (≤ 2 mm) between both groups during the 19:00-19:30 time window and **(E)** during the 08:30-09:00 time window. **(F)** Pre-starved flies spent significantly higher fractions of time on the food patch (inset schematic) as compared to fed flies, with exception of the initial 3 h. * indicate significance levels following pairwise comparison or 2-way ANOVA tests (alpha = 0.05). Horizontal lines indicate significance levels between fed and pre-starved flies from a single group. Curly braces represent identical significance levels between fed and pre-starved from multiple groups. N values are given in brackets.

To summarize, we see that satiation state impacts locomotion and place preference across the day. Within the first hour, starved flies are more active than fed flies, and supposedly compensate their caloric deficit in bouts of short stops until they reach food intake homeostasis. Afterwards, pre-starved flies show reduced activity, particularly during the subjective evening as compared to their fed control group. Interestingly – and despite equivalent hunger motivation – initially starved flies develop increased preference for the food patch over 24 h.

### Initial starvation state impacts movement speed over 24 h

Initially starved flies are equally active as fed flies for the majority of the observed 24 h timespan (Fig. 4A), yet they increasingly confine themselves to the spatially restricted food patch (Fig. 4F). How does this impact the flies’ velocity? As an approximation, we examined the displacement distribution across 24 h between both experimental groups, and found that short displacements of up to 2 mm in the arena were significantly more common in starved flies than in fed flies (Fig. 5A). These short movements are mostly found to be associated with the food patch (movements on patch itself, as well as movements onto patch or off the patch) (Fig. 5B), whereas short movements outside the patch occur with equally low frequency in both experimental groups (Fig. 5C). This suggests that both starved and fed flies have comparable internal drives to move, and the location bias of starved flies towards the food patch is compensated by a significantly higher number of short distance moves of up to 2 mm.

**Figure 5:**
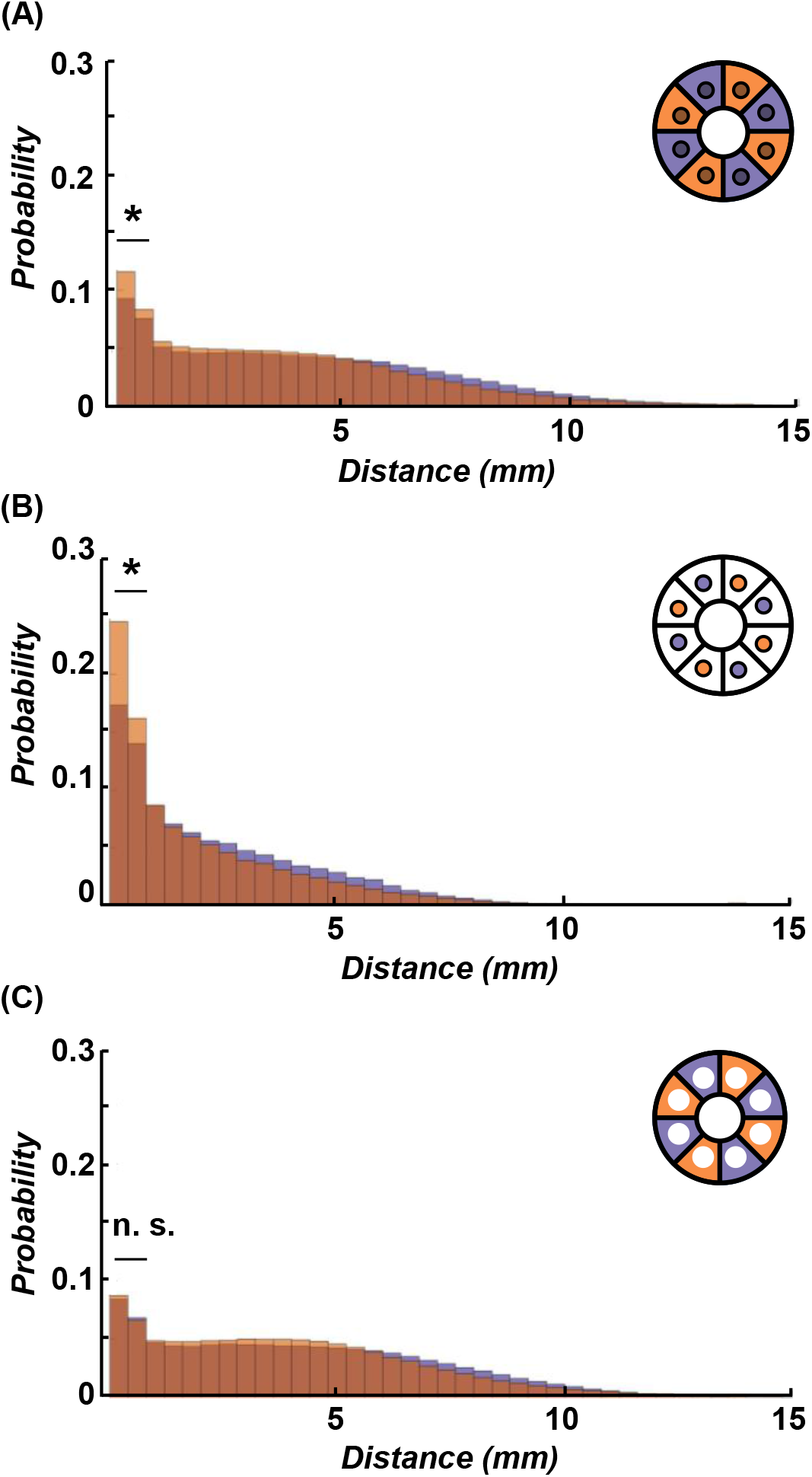
Initially starved flies show more frequent short walks over 24 h. **(A)** Higher probability of short-distance walks (≤ 2 mm) was observed in pre-starved flies (orange) as compared to fed flies (deep purple) within the entire arena (including food patch area, inset schematic) across 24 h. **(B)** The significantly increased short-distance moves for pre-starved flies than those of fed flies were even more pronounced if analysed on the food patch only (inset schematic) **(C)** Probability of flies showing 0.1-2 mm movements were not significantly different between the two groups as observed in the area outside of the food patch (inset schematic). * indicate significance levels following pairwise comparison tests with Bonferroni – Dunn’s correction (alpha = 0.05).

### Non-foraging conditions reinstate evening activity dynamics in initially starved animals

A hungry fly has an intrinsic drive to forage and reach satiety. But how does such a fly behave when we take away the need for foraging altogether, when hunger can be satiated anytime, anywhere? In such a context, we adjusted the assay by removing the food patches and instead lacing all inner surfaces with a homogenous layer of 1 % agarose containing 200 mM sucrose. The raised initial activity in the food patch assay (Fig. 3A) was reduced by about 15 % on omnipresent food in fed flies (Fig. 6A); with about 50 % this effect was even more pronounced in starved flies. The stop lengths in the food-covered arena during the first 30 min are comparable between both experimental groups (Fig. 6B), but there is a robust dichotomy in average speed distribution: starved flies preferably move at average speeds up to 2 mm/s, wherein fed flies travel substantial and significant distances at average speeds between 5-10 mm/s during this time window (Fig. 6C).

**Figure 6:**
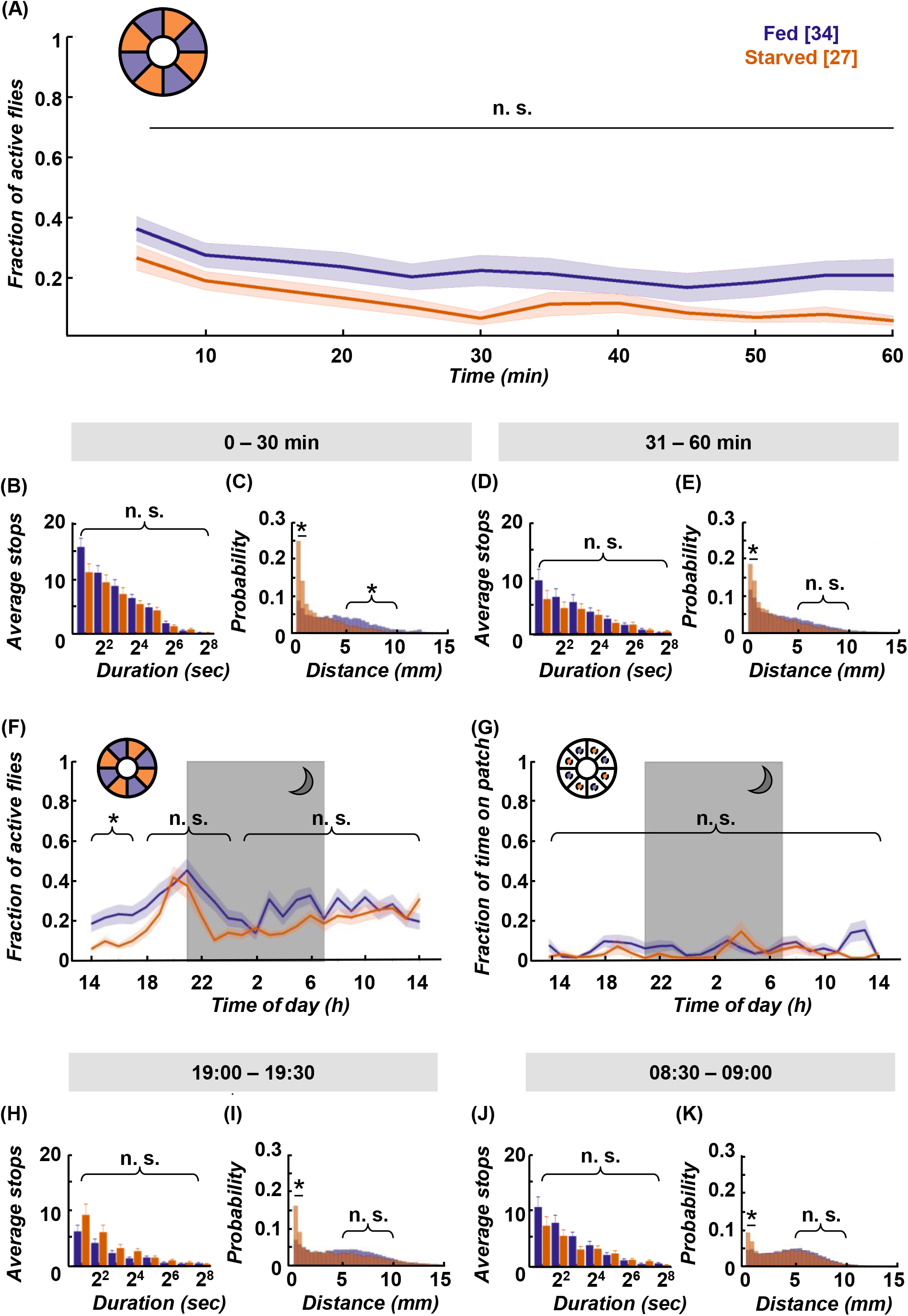
Omnipresent food provision equalises activity levels but not speed. **(A)** Starved (orange, n = 27) and fed flies (deep purple, n = 34) show no significant difference in activity across the first hour on a homogenous sucrose-covered arena (inset schematic), **(B)** together with no significant differences observed for shorter stops (2-7s). X-axis tick labels indicate duration of stops with a log2 scale, where 2^1^ = 2-3s, 2^2^ = 4-7s and so on. **(C)** However, starved flies did show higher frequency of short feeding-related walks (0-2 mm) in the first 30 min than those of fed flies, while fed flies showed a significantly higher frequency of longer distance walks (5-10 mm). **(D)** There was no significant difference observed again in the next 30 min for shorter stops (2-7s) between the two fly groups, but **(E)** pre-starved flies continued to show significantly increased short-distance moves as compared to fed flies. **(F)** Pre-starved flies show comparable activity levels to fed flies, including during the evening activity peak, with no significant difference observed across the remaining 24 h. **(G)** Location probability within virtual food patch-sized areas revealed no significant difference in place preference across 24 h for both pre-starved and fed flies. **(H)** During the 19:00-19:30 time window, both groups showed equal number of short stops (2-7s), with **(I)** consistent higher frequency of short-distance walks seen in pre-starved flies than fed flies. **(J)** No significant difference in short stops (2-7s) was observed during the 08:30-09:00 time window the next morning across both groups, **(K)** and again a significant difference in short-distance moves. Data indicates higher walking speed in fed flies as compared to pre-starved flies. * indicate significance levels following pairwise comparison or 2-way ANOVA tests (alpha = 0.05).

During the subsequent 30 min, when initially starved flies supposedly adjusted their caloric needs, stop lengths remain similar (Fig. 6D). But while starved flies still significantly prefer slow movements, the preference becomes less pronounced and the shift towards higher average velocities reaches comparable levels between 5-10 mm/s in both groups (Fig. 6E). The initially lower activity of pre-starved flies as compared to that of fed flies disappears during the remainder of the 24 h experiment duration. In fact, this even includes a reinstated evening activity peak in pre-starved flies (Fig. 6F). A virtual food patch (i.e. an area of equivalent size to the food patch) in the omnipresent food arena shows no place preferences for both groups (Fig. 6G). It has been reported that flies kept on sugar increase their locomotion in contrast to flies kept on agarose (Lim et al., 2014). Indeed it appears as if overall levels of activity in the omnipresent sucrose arena are slightly elevated, in comparison with fly activity levels in the food patch arena (Fig. 4A, 6F). Taken together, ubiquitous food presentation dampens initial hyperactivity during the first 30 min, and restores the activity peak during subjective evening in starved flies.

### Starved flies maintain reduced walking speed across 24 h on omnipresent food

As mentioned before, homogenous food distribution in the Panopticon results in equivalent stop dynamics for both starved and fed flies. Indeed, this holds true across all examined time windows (Fig. 6B, D, H, J), and is different to what we observed on food patches before (Fig. 3B,C; Fig. 4B,C). Similar to before however, initially starved flies retain a significant prevalence for slower average speeds up to 2 mm/s (Fig. 6C,E,I,K), just as seen in the food patch arena (Fig. 5). This reduced average velocity persists even during later time points when both experimental groups likely have comparable satiety levels - and no carbohydrate restrictions whatsoever. Due to its consistency across assays, the reduced average speed seems to be a conserved feature of the flies’ nutritional history. Taken together, this implies a long-lasting effect of starvation experience on average velocity independent of localised food patches.

## Discussion

The Panopticon represents a cost-effective, Petri dish-based locomotion assay which we utilise in two configurations: i) a classical foraging assay with a single food patch, and ii) an omnipresent food configuration, where every surface in the arena is uniformly covered by substrate. In such a scenario, non-foraging explorative walking behaviour is dissociated from foraging-related search and exploration walking; the flies’ chemosensors are chronically stimulated and every arena location is equally suited to provide food. Due to constant food provision and humidity buffering, longer-lasting recordings (up and beyond 24 h) are possible. One caveat of our experiments is that the flies are exclusively resupplied with a carbohydrate food source. Although the flies’ intake target is heavily skewed towards carbohydrates (Lee et al., 2008; Tatar et al., 2014) and flies can survive on sucrose alone for weeks (Hassett, 1948), we cannot rule out additional effects by ongoing protein deprivation or possible micronutrient shortage. Nonetheless, we only tested male flies which exhibit a more pronounced skew towards sugars than females (Ribeiro and Dickson, 2010). Also, female mating status, egg production and oviposition can affect consumption, as well as food and place choice (Joseph et al., 2009; Carvalho-Santos et al., 2020; Hadjieconomou et al., 2020).

We do not see a pronounced morning activity peak in our assay, as would be expected from circadian studies. However, our experiments are performed in permanent dark, which ensures homogenous illumination and is less disruptive for internal rhythms than constant light (Green, 1964a). But a DD paradigm lacks a distinct visual and thermal Zeitgeber signal for morning onset, which has been shown to be the major influence on the morning activity peak, with little contribution from internal clock genes (Green et al., 2015). Furthermore, individuals within a fly population can exhibit crepuscular, diurnal and nocturnal activity pattern (Pegoraro et al., 2020); the tested flies might be skewed in their allele distribution for such circadian traits. The activity peak during the subjective evening however is very robust. Remarkably, this activity peak is only missing in one condition: in pre-starved flies on the food patch Panopticon. Starved flies do ingest increased amounts of food immediately after food is resupplied (Fig. 3D). Under constant *ad libitum* food conditions, flies tend to not eat to their maximal capacity but rather maintain an almost empty crop (Edgecomb et al., 1994). A full crop cannot only terminate feeding (Gelperin, 1971; Min et al., 2020; Wang et al., 2020a), but also limit post-prandial explorations (Murata et al., 2017). However, it is doubtful that this ‘rest-and-digest’ effect would last very long after the initial voracious re-feeding period. While fed flies always exhibit the evening activity peak, starved flies - which would be expected to have a ‘rest-and-digest’ period after re-feeding - have a reconstituted evening activity peak under omnipresent food conditions (Fig. 6F). This rather indicates a crucial interplay between spatial availability of nutrients and general locomotion motivation across the day to explain the activity differences during the evening in pre-starved flies.

Along with the food patch location preference (Fig. 4F), the following picture emerges: it appears that the motivation for pre-starved flies to sit on the food patch could be not to stray too far from a feeding resource, which outcompetes the motivation for raised activity during the subjective evening. As soon as food is omnipresent, the location preference is gone, and evening activity is reinstated. If the place preference in previously starved flies is also triggered by gustatory activation like local food search (Murata et al., 2017) remains to be shown, for example by providing sweet-only food patches (i.e. arabinose) to rule out caloric involvement. It also remains to be shown if ongoing protein deprivation is involved; however, male flies only seek a protein source after days of prolonged protein starvation (Ribeiro and Dickson, 2010).

The second outcome is less obvious, but robust: independent of food patch presence or omnipresent food, the fed flies move faster than pre-starved flies during the course of the experiments (Fig. 5, Fig. 6C,E,I,K). This could be a delayed effect of the experienced starvation stress; prolonged vibrational stress can lead to reduced voluntary locomotion (Ries et al., 2017). Another possibility could be a long-lasting or even chronic effect on sensory perception (May et al., 2019; Vaziri et al., 2020).

Hunger is known to sensitise certain chemosensory and other circuits at the expense of others, but such sensitisation is usually reversed as soon as the caloric demand is met (Root et al., 2011; Nishimura et al., 2012; Farhan et al., 2013; Inagaki et al., 2014; Longden et al., 2014; Sachse and Beshel, 2016; Grunwald Kadow, 2019; Lin et al., 2019; Wang and Wang, 2019). Habituation and sensitisation of tarsal sugar responses are only described in the minute range (Duerr and Quinn, 1982; Scheiner, 2004; Paranjpe et al., 2012), whereas adaptation of taste-sensing neurons can occur over longer time frames, after chronic dietary intervention (May et al., 2019; Wang et al., 2020b). Exposure to high concentration sucrose might either differentially desensitise gustatory sensilla in fed and pre-starved flies, or the prolonged hunger experience in pre-starved flies leads to a chronic sensitisation of gustatory sensilla. In either case, the threshold to sample the substrate would differ between the two experimental groups, resulting in the observed speed differences in the Panopticon.

The most striking phenotype however is the increasing bias of pre-starved flies towards the food patch (Fig. 4F). A similar difference in food patch interaction (although not on that timescale) has been observed for sitter and Rover alleles of the *foraging* gene; while Rover flies show normal local search behaviour after ingestion, sitter flies tend to stay close to the food source (Pereira and Sokolowski, 1993). Flies are aware of the food patch position within the arena. This is different from learning the spatial arrangement pattern of food patches, which houseflies seem to be not capable of (Fromm and Bell, 1987). But flies can remember locations and learn to efficiently navigate towards previously encountered targets like visual landmarks (Neuser et al., 2008), safe spots (Ofstad et al., 2011), or food sources (Navawongse et al., 2016), even in the dark and without usage of visual or olfactory sense (Kim and Dickinson, 2017).

It appears that the biogenic amine serotonin (5-HT) is involved in such place learning (Sitaraman et al., 2008, 2017; Sitaraman and LaFerriere, 2020). Furthermore, different 5-HT subsets or 5-HT regulation interfere with feeding (Albin et al., 2015; Liu et al., 2015), food seeking behaviour (He et al., 2020), locomotion (Yellman et al., 1997; Howard et al., 2019), sleep architecture (Liu et al., 2019), and quiescence (Pooryasin and Fiala, 2015). Thus, 5-HT manipulation provides a good candidate for further studies in the Panopticon (Tierney, 2020).

Similarly, octopamine (OA) and tyramine (TA) influence locomotion in a state-dependent manner; starvation shifts the OA/TA balance via TBH expression levels and leads to hunger-induced hyperactivity (Yang et al., 2015; Schützler et al., 2019). A single OA neuron signals satiation and stops food-motivated search (Sayin et al., 2019), and the same neuron can initiate feeding behaviour (Youn et al., 2018). Also, OA influences AKH signalling for diurnal pattern generation (Pauls et al., 2020), and might be affected in starved flies during the blocked evening activity peak. Given the pleiotropic actions of OA, place preference may be impacted as well (Selcho and Pauls, 2019).

The food patch location is most likely associated with food reward in both fed and starved flies, analogous to odour associations with caloric value (Burke and Waddell, 2011; Fujita and Tanimura, 2011; Huetteroth et al., 2015; Ichinose et al., 2015; Musso et al., 2015; Yamagata et al., 2015; Zhang et al., 2015). Such long-lasting, food-related odour memories are stored in the mushroom body (MB) (Krashes and Waddell, 2008), and it is clear that this structure, especially its zonal dopaminergic modulatory innervation, has a central instructional role in motivational foraging and feeding (Tsao et al., 2018; Musso et al., 2019; May et al., 2020).

In the Panopticon experiment, the food-place association could be enforced by two factors in starved flies: First, the absolute amount of food that is ingested by starved flies within the first 30 min is bigger, since they supposedly need to cover their caloric deficit (Fig. 3D). Secondly, the lack of caloric signals during starvation renders food-associative MB circuits particularly sensitive to the next food encounter (Hirano and Saitoe, 2013; Hirano et al., 2013, 2016; Plaçais et al., 2017; Wu et al., 2018). For example, starved flies, contrary to fed flies, do not require additional sleep to consolidate a food-odour association (Chouhan et al., 2020). It appears the subjectively perceived value of food is higher in starved flies than in fed flies. So during this time of transition in a new environment, both quantity and perceived quality of the ingested food on the food patch would be higher for starved flies. These two effects together could lead to a strong and long-lasting positive association with the food patch location that influences location decision making beyond nutritional demand for the subsequent 24 h.

However, some issues are not addressed by this explanation. Retrieval of a food-associated memory depends on the motivational incentive of hunger; a starved fly will utilize an olfactory food association to increase its chances to feed, whereas fed flies will only do so after being starved once more (Krashes and Waddell, 2008). Similarly, starved flies would have a higher incentive to retrieve and use their place memory of the food patch location, and indeed, starved flies exhibit a higher and more frequent return rate to a known food patch, be it real or virtual (Corrales-Carvajal et al., 2016; Kim and Dickinson, 2017; Murata et al., 2017; Corfas et al., 2019; Haberkern et al., 2019).

In this regard it is unlikely that the lasting food patch preference in the Panopticon depends on concurrent hunger as the motivational drive to retrieve spatial memory. All other food-related behaviours like activity, stop distribution or sip number are aligned between starved and fed flies within an hour, indicating comparable hunger motivation from then onwards (Fig. 3), and protein hunger only starts to influence male food choice much later (Ribeiro and Dickson, 2010). It also needs to be taken into account that food association in the Panopticon is not formed with odours but with place, and associations of spatial features involve the central complex (Liu et al., 2006; Stern et al., 2019); it is equally possible that plastic changes in this structure contribute to the observed location preference. In a place learning assay, the unavoidability of an aversive heat stimulus can boost its reinforcing propensities (Sitaraman and Zars, 2010); it is feasible that the same is the case for unavoidable starvation.

How could this work? This reminds of long-lasting, non-associative and associative learning effects, mostly known from the classic *Aplysia* gill- and siphon-withdrawal reflex (Pinsker et al., 1973; Cassel, 2010; Cai et al., 2011). Repeated noxious stimuli on the tail or neck of this sea slug sensitise the animal so that subsequent normal mechanosensory stimulation of the siphon results in exaggeratedly strong gill and siphon withdrawal (Frost et al., 1985). The same pathways were shown to be also involved in classical conditioning experiments of the gill- and siphon-withdrawal reflex (Hawkins et al., 1983). In our case, the sensitising noxious signal is replaced by an internal need (hunger), whereas the sensitised motor response is replaced by a decision-making circuit (bias to stay on food).

But why does this effect appear to become stronger over time? It might be possible that the starved flies generalise from the starvation experience, and hence seek proximity to the food source. Increasing generalisation of an aversive stimulus over time is not only observed for odour-shock learning in flies (König et al., 2017), but is also a characteristic feature of post-traumatic stress disorders in both animals and humans (Stam, 2007). A similar long-lasting effect has been described for predator-induced oviposition preference (Kacsoh et al., 2015). Here, gravid female flies are exposed to parasitoid wasps for several hours. After removing the wasps, the females choose ethanol-laced patches over control patches for days (fly larvae have a higher ethanol tolerance than wasp larvae). As in the Panopticon, prolonged exposure to a distressing stimulus (hunger or parasitoid wasps) influences a later choice (place preference) even after the stressor was removed. Interestingly, MB inhibition and several memory mutants abolished this long-lasting skew exclusively after wasp removal, but not under immediate threat; it will be interesting to see how MB function and memory genes impact the place preference on the Panopticon.

## Outlook

We present here a new paradigm to examine locomotion behavior and place preference, under foraging conditions (food patch Panopticon) or under chronic chemosensory stimulation (omnipresent food Panopticon). The food patch Panopticon will help to examine the neuronal circuits underlying long-lasting effects of starvation on place preference, and how this apparently non-associative process relates to known associative long-lasting memory function.

In the omnipresent food Panopticon, we will be able to assess the influence of state-modulating or state-mediating substances like biogenic amines or neuropeptides and their receptors on locomotion parameters, without interference of foraging-motivated movement. Being able to do this over prolonged periods will help to discern long-lasting pleiotropic effects of these effectors (Martelli et al., 2017; Dreyer et al., 2019; Nässel et al., 2019).

## Data availability statement

The 3D .stl-file of the Panopticon and the MATLAB scripts are available on Github under this link: https://github.com/dmahishi/Panopticon.git; the datasets for this study and further details on the Panopticon are available from the authors upon request.

## Author contributions

DM, TT, and WH conceived and designed the experiments. DM and RH performed the experiments. DM and TT wrote the MATLAB script. DM, TT and WH analysed the results and wrote the article. All authors provided comments and approved the manuscript.

## Acknowledgments

We thank Ingo Kannetzky for expert help with 3D printing and design. We also thank Andreas Thum, Bert Klagges and Kathrin Steck for their support and input on all levels.

## Supplementary Material

1. 3D-printable .stl file of Panopticon
2. FIJI-Macro for tracking
3. MATLAB code & Documentation

---

**Figure S1:**
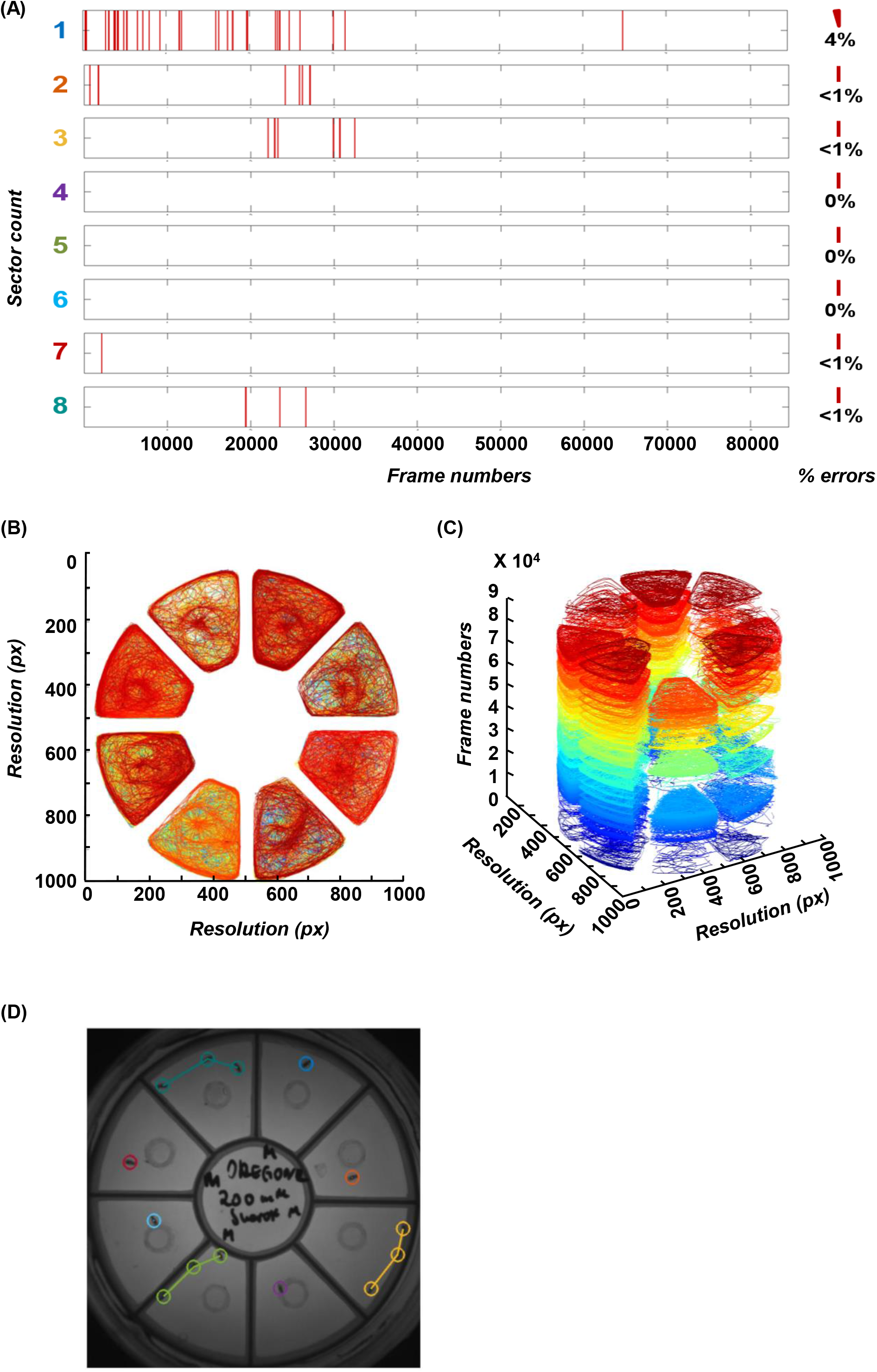
Representative plots for multi-step quality check. **(A)** Temporal distribution of failed/missed detections with total error rate (%) indicated on the right panel for individual flies/sectors across 24 h. **(B)** Walking path traces for the entire arena divided by sectors, with **(C)** 3D representation, expandable on the Z-axis representing walking traces across the total duration of activity recording for 24 h. Time scales represented as frame numbers (@ 1 frame/sec). Blue-yellow-red colour transition represents walking traces from 0-n frame count (across total duration of the experiment). **(D)** To visually identify long-distance movements associated with potentially false-positive detections, corresponding images for top ten frames with flies covering the longest distances can be extracted (example image for Sector 8). Such images include up to 2 consecutive long distance moves as indicated by individually coloured walking trace lines and the fly itself encircled with the respective colour. Apart from the queried sector, all the other sectors also indicate distances moves in corresponding frames.

